# Rational engineering of industrial *S. cerevisiae*: towards xylitol production from sugarcane bagasse

**DOI:** 10.1101/2021.12.08.471450

**Authors:** Fellipe da Silveira Bezerra de Mello, Carla Maneira, Frank Uriel Suarez Lizarazo, Sheila Nagamatsu, Beatriz Vargas, Carla Vieira, Thais Secches, Alessando L V Coradini, Maria Augusta de Carvalho Silvello, Rosana Goldbeck, Gonçalo Amarante Guimarães Pereira, Gleidson Silva Teixeira

**Author notes:** These authors contributed equally to this work (Fellipe da Silveira Bezerra de Mello, Carla Maneira).

## Abstract

**BACKGROUND:** Sugarcane hemicellulosic material is a compelling source of usually neglected xylose that could figure as feedstock to produce chemical building blocks of high economic value, such as xylitol. In this context, *Saccharomyces cerevisiae* strains typically used in the Brazilian bioethanol industry are a robust chassis for genetic engineering, given their robustness towards harsh operational conditions and outstanding fermentation performance. Nevertheless, there are no reports on the use of these strains for xylitol production using sugarcane hydrolysate.

**RESULTS:** Potential single-guided RNA off-targets were analyzed in two preeminent industrial strains (PE-2 and SA-1), providing a database of 5’-NGG 20 nt sequences, and guidelines for the fast and cost-effective CRISPR-editing of such strains. After genomic integration of a NADPH-preferring xylose reductase (XR), FMYX (SA-1 *ho*Δ::*xyl1*) and CENPKX (CEN.PK-122 *ho*Δ::*xyl1*) were tested in varying cultivation conditions for xylitol productivity to infer influence of the genetic background. Near-theoretical yields were achieved for all strains, however the industrial consistently outperformed the laboratory strain. Batch fermentation of raw sugarcane bagasse hydrolysate with remaining solid particles represented a challenge for xylose metabolization and 3.65 ± 0.16 g/L xylitol titre was achieved by FMYX. Finally, quantification of NADPH - cofactor implied in XR activity - revealed that FMYX has 33% more available cofactors than CENPKX.

**CONCLUSIONS:** Although widely used in several *S. cerevisiae* strains, this is the first report of CRISPR-Cas9 editing major yeast of the Brazilian bioethanol industry. Fermentative assays of xylose consumption revealed that NADPH availability is closely related to mutant strains’ performance. We also pioneer the use of sugarcane bagasse as a substrate for xylitol production. Finally, we demonstrate how industrial background SA-1 is a compelling chassis for the second-generation industry, given its inhibitor tolerance and better redox environment that may favor production of reduced sugars.

## BACKGROUND

The recent advancements in industrial biotechnology have enabled the effective reuse of agro-industrial residues in a range of applications. Sugarcane bagasse, for instance, is the major by-product of the sugarcane industry - a billionaire market that employs millions of people worldwide [1]. The abundance of this lignocellulosic material makes it an ideal substrate for exploring the potentials of underutilized pentose sugars. Recently, the production of second-generation bioethanol (E2G) from xylose-rich waste raised awareness on the potential holistic utilization of this biomass [2]. However, many research groups have lately assessed that the generation of alternative products, such as xylitol, might stand out as an even more promising alternative to compensate for the costs of fuel production from sugarcane substrates [3,4].

Xylitol is a five-carbon sugar alcohol that occurs naturally in certain fruits and vegetables [5]. It presents sweetness equivalent to sucrose while having just 60% of its calorie content, being mostly used as a natural sweetener in chewing gums [6]. Besides its well-established application in the food and beverage industry [7], xylitol has great potential in the pharmaceutical industry [2,8,9]. Altogether, xylitol properties speak for themselves when it comes to understanding why its demand is expected to grow in a market with increasingly health- and weight-conscious consumers. By 2025, xylitol market is expected to reach $1.37 thousand million, with a price range of $4000-5000 per tonne [10].

Biotechnological routes have been considered a relevant substitute to the conventional chemical method of xylitol production, as they can be based on a mixture of sugars and save on energy and substrate purification costs [2]. There are reports of microbial processes based on bacteria, fungi, and yeast for xylitol production, being the last considered the best producers [11]. During these bio-based processes, D-xylose is reduced in a single step into xylitol by the NAD(P)H-dependent enzyme Xylose Reductase (XR), which is further secreted [12]. In order to enable high titre xylitol production in *S. cerevisiae*, the heterologous expression of *Scheffersomyces stipitis’* XR-encoding *XYL1* gene or overexpression of endogenous *GRE3* are common strategies that have been widely applied [13,14].

While xylitol production by genetically modified *S. cerevisiae* using non-detoxified hemicellulosic hydrolysates from corncob and rice straw has been reported [15–18], there is no register on the inquiry of sugarcane bagasse as substrate. In this context, the use of industrial yeast strains adapted to commercial fermentation, - especially the ones applied in the sugarcane-to-ethanol industry - stands out as important chassis towards enabling xylitol production in recalcitrant conditions [19]. Examples of top-performing indigenous *S. cerevisiae* encompass the Brazilian bioethanol strains Pedra-2 (PE-2) and SA-1, that present outstanding fermentation capacity although subjected to numerous stresses [20]. Prospects on the use of these strains targeting the E2G industry have already been described: PE-2 has been explored for xylose consumption [21] and strain SA-1 was recently reported as highly resistant to major aldehyde inhibitors found in the sugarcane hydrolysate - such as 5-hydroxymethylfurfural (HMF) and furfural [22]. These pieces of evidence indicate them as compelling chassis for xylitol production using sugarcane waste biomass.

In order to obtain relevant recombinant industrial strains for xylitol production, their rational genetic engineering deems necessary. Even though the CRISPR-Cas9 system is widely applied in *S. cerevisiae*, usage parameters are often strain-dependent and require fine adjusting [23]. PE-2 and SA-1 have a very specific genetic background, contrasting to the S288c model that has set the CRISPR-editing systems utilized in this microorganism. Besides presenting higher ploidy, these industrial strains were conditioned to specific environmental adaptations that resulted in a highly heterozygous genome [24], distinctive from the common haploid laboratory strains. Although very relevant, there are no reports of successful CRISPR-editing of Brazilian yeast strains of economic significance to the ethanol business.

Here, we assess the efficiency of xylitol production of engineered bioethanol *S. cerevisiae* strains after establishing optimal CRISPR-Cas9 editing parameters. We cover work towards mapping differences between single-guided RNA (sgRNA) sequences in PE-2 and SA-1 in relation to laboratory S288c, providing a sgRNAs database for these strains. We also set an efficient CRISPR-based genomic editing protocol for the working strains. Following, cultivation of an engineered SA-1 in rich media resulted in total xylose metabolization, and nearly theoretical xylitol yield was achieved. Regarding xylitol productivity in sugarcane bagasse hydrolysate, recombinant SA-1 was able to produce 3.65 ± 0.16 g/L of the reduced sugar, outperforming mutant CENPK-122. NADPH quantification revealed that the industrial background has 33% more cofactor availability than the laboratory, gathering evidence that aldehyde-resistant SA-1 has a favorable redox environment that can improve XR activity and serve as an interesting chassis for the second-generation industry.

## MATERIALS AND METHODS

### STRAINS, PLASMIDS AND MEDIA

All *S. cerevisiae* strains and plasmids used in this study are described in **Table 1**. Yeasts were cultivated in YPD medium (10 g/L yeast extract, 20 g/L peptone, 20 g/L glucose) for inoculum and propagation purposes, or YPDX (10 g/L yeast extract, 20 g/L peptone, 20 g/L xylose and varying glucose concentration) for xylitol production assays. Cultivation was carried out at 30 °C and 250 rpm, unless otherwise noticed. Assays were performed aerobically (80 mL medium in unsealed 250 mL Erlenmeyer flask) or semi-anaerobically (80 mL medium in rubber stopper-sealed 250 mL Erlenmeyer flask). Geneticin (200 μg/mL g418) was used for the selection of yeast transformants carrying a KanMX marker, present in plasmid pGS. *Escherichia coli* DH5□ was used for propagation and storage of vectors and was cultivated in Luria-Bertani (LB) broth (10 g/L tryptone, 5 g/L yeast extract, 10 g/L NaCl) at 37 °C and 250 rpm, when in liquid media. Ampicillin (amp, 100 μg/mL) was added for the selection of bacteria colonies expressing plasmids. All media previously described were added 15 g/L of agar for solidification. Microorganisms stock solution were kept at −80 °C in media containing 25% glycerol, for long term storage.

**Table 1:**
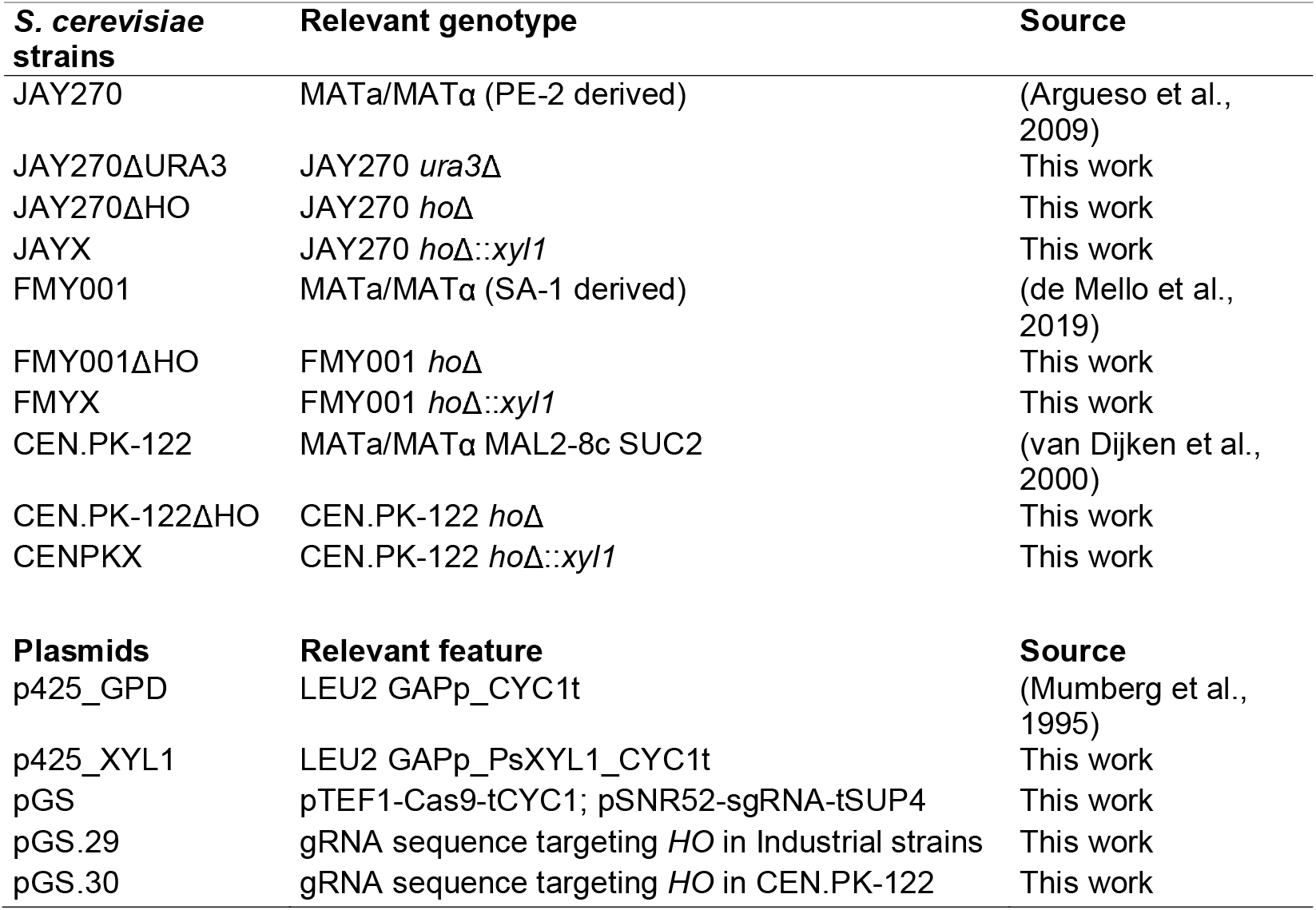
Main *S. cerevisiae* strains and plasmids used in this work.

### PE-2 AND SA-1 sgRNA OFF-TARGET ANALYSIS

All NGG PAM sequences (NoNAG) CRISPR-Cas9 targets in the yeast genome, previously disclosed by DiCarlo *et al*. [25] were compared to the publicly available genomes of both JAY291 [24] and FMY097 (SA-1 derived) [26]. Further, off-target possibility was defined according to the recurrence of sgRNAs - multiple identical sequences or bearing up to 3 single nucleotide polymorphisms - using bowtie (version 1.0.0) (Langmead et al., 2009) set with parameters -l 12 -v 3 -a. A homemade script in Perl was developed to count the single polymorphisms differences identified in the alignment output (sam file) for each reference (JAY291 and FMY097) and sgRNA.

### sgRNAS SYNTHESIS AND ASSEMBLY

sgRNA sequences were predicted using CHOPCHOP [27] and the sequences were compared to the off-target analysis previously performed. A ligation-based cloning approach similar to that described by Laughery *et al*. (2015) [28] was performed. In summary, 300 μM of complementary primers containing 20 bp overlap - target sequence - and 4 bp of homology to the expression vector were hybridized in a reaction containing 10 μL of 10x T4-ligase buffer (New England Biolabs). The mixture was submitted to 70 decreasing cycles of a minute each - touchdown - from 95 to 25°C and then connected to the previously linearized vector through a reaction containing 15 ng of the previous hybridization, 50 ng linearized plasmid, 2 μL of 10x T4-ligase buffer, and 1 U of T4-ligase (New England Biolabs). After ligation, each reaction was dialyzed and transformed into *E. coli*. After transformation, positive clones were selected on LB-amp. The sgRNA insertion was confirmed by PCR and sequencing.

### DONOR DNA SYNTHESIS

Knock-out (KO) double-stranded donor DNAs (dsOligos) were synthesized using 55 bp DNA primers containing 20 overlapping nucleotides. The synthesis was carried out in a template-free Phusion High-Fidelity DNA Polymerase (ThermoFisher) reaction. Each reaction was kept in an appropriate thermocycler for 5-10 cycles of denaturation at 98 °C, annealing at 50 °C and extension for 15 seconds. The synthesis product - 90 bp dsOligo - was confirmed by gel electrophoresis using 2.5% agarose gel. Knock-in donors were amplified from genomic DNA. The homology regions for subsequent gene repair were either based on primer extensions (for homology regions up to 60 bp) or on previously amplified regions flanking the to-be-edited area.

### CLONING AND GENERAL MOLECULAR BIOLOGY

The *XYL1* gene from *Sc. stipitis* was cloned in the *BamH*I-digested p425-GPD [29] through gap repair in *S. cerevisiae*. The p425-XYL1 plasmid was used as the replication material for the XR donor cassette. All primers used in this study are described in the **Supplementary Material (Table S1)**. Plasmids propagated in *E. coli* were extracted using the alkaline lysis procedure described by Birnboim and Doly (1979) [30]. Yeast genomic DNA was extracted using the standard Phenol-Chloroform protocol [31]. Polymerase chain reactions (PCR) were carried out with Phusion High-Fidelity DNA Polymerase (ThermoFisher) according to the manufacturer’s instructions. Yeast transformation was performed using the LiAc/SS carrier DNA/PEG method, described by Gietz *et al*. (2007) [32]. Vectors were inserted in *E. Coli* using the traditional electroporation protocol [33].

### HO *locus* NEUTRALITY TEST

To confirm the neutrality of the *HO* gene KO genotype in the fitness of strains FMY001 *ΔHO*, CEN.PK-122 *ΔHO* and JAY270 *ΔHO*, their phenotype was assessed by biomass yield after growth in optimal conditions. In duplicates, cultivation of these strains in 20 mL of YPD was performed at 30 °C and 250 rpm of orbital shaking in unsealed 125 mL Erlenmeyer (aerobic growth). Optical density (OD_600nm_) of the culture was measured with a spectrophotometer after 12 hours of growth.

### XYLITOL PRODUCTION ASSAYS IN YPDX

Transformants were grown aerobically in YPD medium in Erlenmeyer flasks overnight for inoculum. The saturated culture was centrifuged at 2000*g* for 5 minutes and the cells were washed with sterile distilled water. The pellet was resuspended in order to achieve a fermentation initial OD_600nm_ of 0.5 or 1.0. For the initial assays concerning xylitol productivity, strains FMYX, CENPKX and JAYX were cultivated in YPDX (5 g/L glucose), initial OD_600nm_ 0.5, semi-anaerobic growth for 120h. HMF influence over xylitol productivity of strains FMYX and CENPKX was assessed with YPDX (5 g/L glucose) medium supplemented with 0.5 or 2 g/L HMF, in the same cultivation conditions previously mentioned. Analysis of xylitol productivity in varying cultivation conditions was carried out with strains FMYX and CENPKX in different settings: initial OD_600nm_ of 0.5 or 1.0; glucose concentration of 10, 20 or 30 g/L; aerobically or semi-anaerobically for 104h. All cultivation assays were carried out in triplicates. Samples were withdrawn for HPLC analysis.

### NADPH/NADP QUANTIFICATION

NADP/NADPH concentration was measured using the NADP/NADPH-Glo (Promega) assay kit. All the *S. cerevisiae* strains used in this experiment were grown in YPD and harvested at the mid-exponential growth phase (OD_600nm_ 0.8 - 1.0). 1 mL of OD_600nm_ 1.0 of each strain was centrifuged and resuspended in 150 μL of the NADP(H) extraction buffer. Further procedures followed the manufacturer’s protocol.

### XYLITOL PRODUCTION IN SUGARCANE HYDROLYSATE

Steam-exploded sugarcane straw (23.9% solids) was donated by GranBio SA (Bioflex plant) and used for hydrolysate production. Operational conditions and chemical composition of the pretreated material were not disclosed. Cellic CTec3 (6% w/w glucan) was used for enzymatic hydrolysis, which occurred in 250 mL Erlenmeyer flasks containing 17g of pretreated sugarcane straw (dry basis) and distilled water to complete 100g total working volume. Ammonium hydroxide was used to adjust the pH to 5. Hydrolysis reaction occurred under 250 rpm agitation, at 50 °C, for 72 h. The same 250 mL Erlenmeyer flasks used for the enzymatic hydrolysis (100g reaction), added by 4 ppm ampicillin, were used for fermentation. Fermentation occurred for 120 hours with initial OD_600nm_ 1, under the same conditions used for inoculum preparation (aerobic growth at 30 °C and 150 rpm). Samples were withdrawn for HPLC analysis. All fermentations were conducted in triplicate.

### ANALYTICAL METHODS

Glucose, xylose, xylitol, glycerol, acetic and formic acid, ethanol, furfural, and HMF concentrations were determined by high performance liquid chromatography (HPLC, Allience HT, Waters, USA). Compounds separation occurred in a Bio-Rad HPX-87H column at 35 °C, using 5 mM sulfuric acid as mobile phase, at 0.6 mL/min. A photodiode array detector at 280 nm was used for furfural and HMF; a refractive index detector at 35 °C was used for the other compounds.

### FERMENTATION PARAMETERS CALCULATION AND STATISTICS

Fermentation parameters were defined as follows: xylose conversion (%) as the ratio of the amount of xylose consumed (g/L) by the initial xylose loading (g/L); xylitol yield (g/g) as the amount of xylitol produced (g/L) per consumed xylose (g/L); xylitol productivity (g/L.h) as the amount of xylitol produced (g/L) divided by fermentation time (h). All data are presented as mean ± standard deviation. Statistical analysis between means was assessed with Tukey’s test (0.95 confidence interval), calculated using Minitab version 17 (Minitab Inc., State College, PA, USA).

## RESULTS

### Efficient sgRNA design based on specific genome information is key for editing industrially relevant strains

With the consolidation of CRISPR-Cas9 as one of the most effective DNA editing tools in *S cerevisiae*, precise sgRNA design for the endonuclease activity is often the foremost step when one intends to modify genomic sequences. A primary effort into the rational genetic engineering of wild type strains - such as bioethanol yeasts PE-2 and SA-1 [20] - is to consider how their genomes differ from the S288c model. As most of the trustworthy sgRNA prediction analysis software relies on the model yeast S288c for establishing its efficiency prognosis, the degree of accuracy of these programs can be limited in strains with diverging backgrounds. For instance, single nucleotide polymorphisms’ (SNP) presence in PE-2 is calculated to be as high as 65,000 when analyzing the whole genome in comparison to S288c [24]. Therefore, a complete analysis of all possible sgRNAs encompassed in PE-2 and SA-1 genomes and their mismatch probability in relation to S288c sgRNAs was performed.

In summary, all CRISPR-Cas9 NoNAG targets in the yeast genome, previously disclosed by DiCarlo *et al*. (2013) [25] were compared to the publicly-available genomes for both the JAY291 strain [24] - PE-2 haploid derived from the JAY270 diploid - and the FMY097 strain [22,26] - haploid derived from the FMY001 strain – SA-1 diploid-derived. The analysis detected sgRNAs with identical hits in the industrial strains’ genomes, as well as highly similar sgRNAs that could lead to off-target activity of the Cas9 enzyme. SgRNAs were considered to lead to off-targets due to either (1) the existence of another fully identical 20bp DNA sequence 5’ to a PAM in the genome, (2) the presence of highly similar sequences (up to 3 bp differentiating them), (3) the nonexistence of the original sgRNA sequence found in the S288c analysis or (4) a combination of these events. The full list of potential sgRNAs and their off-target probabilities in PE-2 and SA-1 in relation to all S288c NoNAGs has been provided in .csv files (**See Supplementary Files, Tables S2 and S3**). **Figure 1** shows a graphical overview of the results obtained.

**Figure 1:**
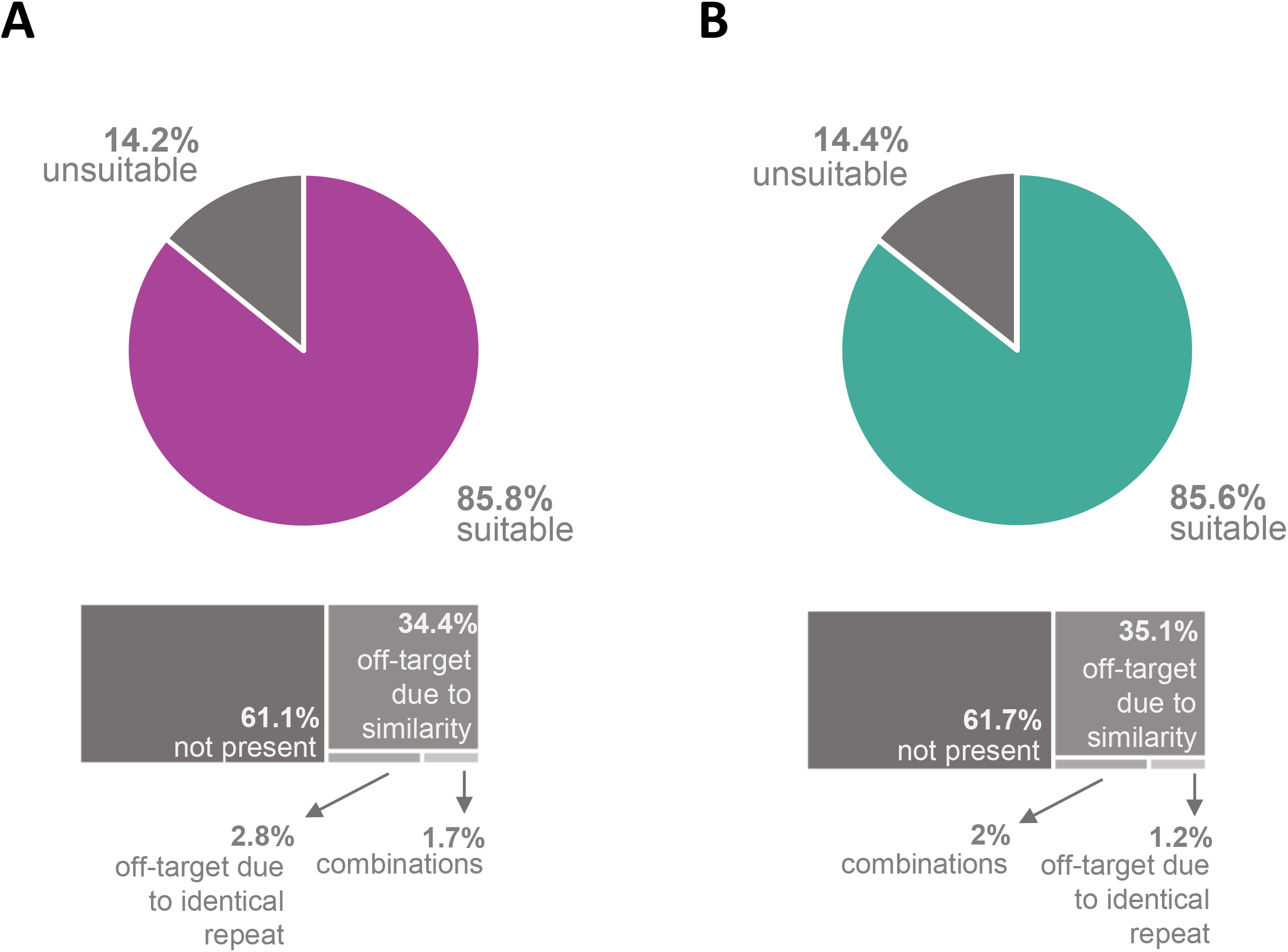
Single-guided RNA (sgRNA) prospection in bioethanol *S. cerevisiae* strains. Sequences predicted for model yeast S288c were used for comparison. **A.** sgRNA suitability for JAY291 (PE-2 segregant). **B.** sgRNA suitability for FMY097 (SA-1 segregant).

For PE-2, out of the 524,288 sgRNAs predicted for the s288c model, 511,433 found hits in the PE-2 genome, 85.8% of them were considered suitable, while the other 14.2% were classified as ineligible due to one of the four off-target criteria. As for SA-1, there were 513,159 hits in the genome: 85.6% were considered suitable, while 14.4% were disqualified. In both analyses, sequences were mainly classified as unsuitable due to nonexistence of the original sgRNA sequence, followed by off-target due to similarity (up to 3 mismatches).

### Synthesis of CRISPR-Cas9 editing components for Brazilian bioethanol strains

Following genome-wide sgRNA screening in industrial strains PE-2 and SA-1, we proceeded to establish a simple and cost-effective yeast transformation toolkit for our strains. When it comes to the CRISPR-Cas9 system delivery, an approach similar to that described by Stovicek *et al*. (2015) [23] was chosen. However, a single-plasmid (named pGS) plus double-stranded oligo system was applied to simplify the process. pGS bears both the Cas9 gene under the strong TEF1 promoter, and the sgRNA spacer 5’ to the SNR52 promoter. Additionally, for the easy cloning of sgRNAs into the plasmid, a single-enzyme-based restriction and ligation strategy was implemented. The complete strategy is detailed in **Material and Methods** (**see Supplementary Figure S1A**).

### Effective CRISPR-Cas9 editing in the Brazilian industrial yeast strains

In order to establish optimal transformation conditions for our strains, we selected the JAY270 strain (PE-2 isolate) as a model yeast for this assay. The KO of the *URA3 locus* was chosen for this assessment, due to the ease in screening for edited colonies based on auxotrophy. In short, JAY270 was subjected to 15 different transformation conditions bearing variable concentrations of the Cas9-sgRNA pGS vector and the donor DNA - a 90 bp dsOligo with a stop codon replacing the PAM. The sgRNA sequence used for this purpose was validated using the information provided in the first result section. As depicted in **Figure 2A**, the editing efficiency was over 95% for 4 out of the 15 tested conditions. The condition bearing 1500 ng of the vector and 1000 ng of the donor DNA stands out with a 99% efficiency rate (**Figure 2A**). This condition was classified as optimal and used in all subsequent transformation events for both the PE-2 and SA-1 strains.

**Figure 2:**
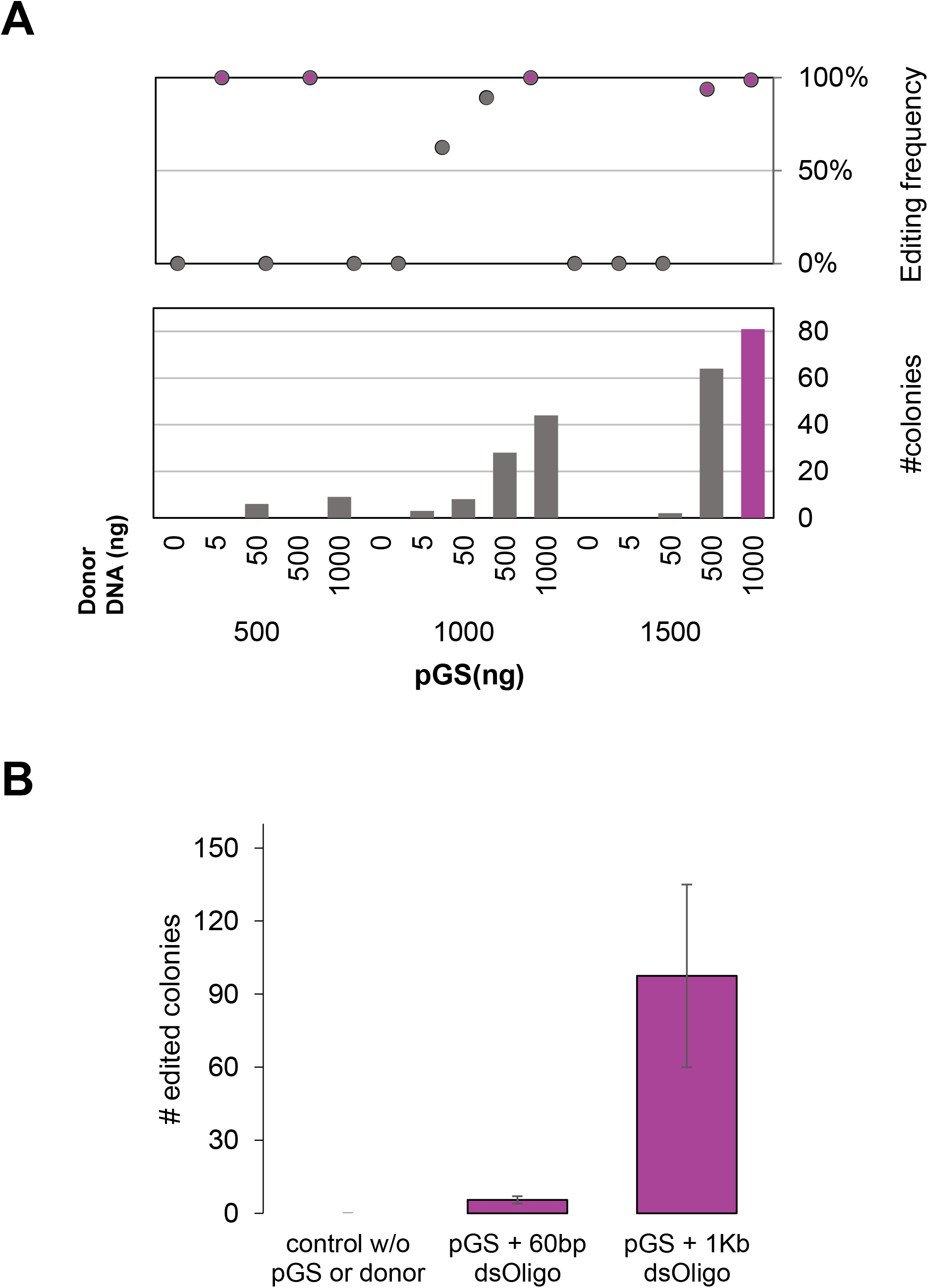
Standardization of a CRISPR-Cas9 system for the genome editing of diploid industrial strains with plasmid pGS (Cas9 + sgRNA scaffold), using JAY270 (PE-2 derived) as a proof of concept. **A.** Editing efficiency of the *URA3 locus* knock-out using different concentrations of plasmid pGS and a 90bp-long donor DNA. **B.** Editing efficiency of the *URA3 locus* knock-in using different donor overhang homologies to the editing site.

After confirming the system functionality for KOs, we then assessed efficiency for whole-gene integration. The JAY270ΔURA3 strain was used as a background for a gene knock-in transformation assay. The optimal transformation condition (1.5 μg pGS and 1 μg donor DNA) was applied for re-integration of the *URA3* gene into the previously disrupted *URA3 locus*. Additionally, two donor DNA homology-overhang with varying lengths were tested. As already observed in previous works, a correlation between editing efficiency and the length of donor DNA overhang homologies is shown in **Figure 2B**.

Due to the high number of edited colonies (on average 97.5) for the donor DNA bearing 1 Kb-long homologies this condition was considered excellent. However, even low-cost options such as the use of the donor DNA bearing 60 bp-long homologies were considered pertinent (on average yielded 6 edited colonies). Overall, the knock-in assay using pGS and previously described transformation conditions led to successfully edited colonies, proving the efficiency of the CRISPR-Cas9 system and transformation method for gene integration for the PE-2 strain.

In a similar fashion, we tested the efficiency of the system here described to edit haploid cells. LVY34.4 [21], a haploid derivative of the PE-2 strain, was subjected to this methodology for the KO of the *LEU2 locus*. Again, high editing efficiency rates very close to 100%, as well as a high number of edited colonies (92 out of 93 successfully transformed colonies) were obtained (**See Supplementary Figure S2**).

### XYL1 integration in diploid industrial strains

Following, we sought to apply the herein developed CRISPR-based genome editing toolkit to endow widely used bioethanol strains JAY270 (PE-2) and FMY001 (SA-1) with higher xylitol production performance. For that, the integration of a *SsXYL1* expression cassette was designed in the *HO locus* (HOmothalic switching endonuclease). The top-indicated sgRNA for editing the *HO* allele enclosed a SNP - corresponding to nucleotide 789 in the gene ORF - in the industrial PE-2 and SA-1 strains when compared to the model S288c. In fact, sequencing of both strains uncovered that diploid SA-1 is heterozygous for this mutation, while PE-2 is homozygous. Plasmids pGS.29 and pGS.30 were, therefore, designed to contain sgRNAs targeting each variation of the *HO locus*. The plasmids contained a single base-pair mutation in the sgRNA sequence differentiating between them (**see Supplementary Table S1**), allowing the specific editing of PE-2 or CEN.PK-122, respectively. Importantly, the diploid laboratory strain CEN.PK-122 [34] - harboring the same *HO* sequence as in S288c - was selected as a control to guarantee a proper evaluation of the industrial background influence in xylitol production, explored in the next sections.

Effective sgRNA design was assessed in transformations containing solely the pGS plasmid (no donor DNA co-transformed). As Cas9-induced double strand break is preferably repaired by homology-directed mechanisms in *S. cerevisiae* [35], a high incidence of cell-death - leading to an easy-to-spot low colony count pattern - is achieved when no donor DNA is co-transformed with an efficient sgRNA-containing plasmid. Being so, a transformation of both industrial strains and the control with pGS.29 and pGS.30 alone, revealed that the single base-pair change was crucial for the Cas9 endonuclease activity - *i.e*., pGS.29 successfully edited PE-2 and not CEN.PK-122, while pGS.30 only worked for the latter. Both plasmids did not present activity on SA-1, given its heterozygous *locus* (**See Supplementary Figure S3**). Being so, SA-1 transformation was carried out separately in segregant haploids FMY034 (MAT□) and FMY097 (MATa) [22] - both with the same SNP in gene *HO* as in PE-2 - and further crossed.

Following the confirmation of effective sgRNAs for all strains, we investigated the *HO* KO neutrality in the industrial strains’ fitness, given that no information regarding this genotype is available for such yeast background. A non-coding cassette, pGAP-tCYC1, was integrated into SA-1 and PE-2’s gene using pGS.29. Growth of *HO* knocked-out strains in optimal conditions certified that this allele interruption does not affect cell cultivation for these strains (**See Supplementary Figure S4**).

Next, the *XYL1* expression cassette (pGAP-SsXYL1-tCYC1) was integrated into CEN.PK-122 and PE-2 genomes, with 40 bp flanking homology, leading to 11% and 28% editing efficiency, respectively. Subsequently, the donor cassette was amplified from the transformed PE-2 with 300 bp overhang homology to the editing site and used in the transformation of SA-1 segregants, resulting in 83% editing efficiency (**See Supplementary Figure S5**). Edited strains were labeled CENPKX (CEN.PK-122 *ho*Δ::*xyl1*), JAYX (PE-2 *ho*Δ::*xyl1*) and FMYX (FMY034/FMY097 *ho*Δ::*xyl1*).

### Xylitol production by modified bioethanol Brazilian strains: a prospect

In this section, we explore how edited strains perform regarding xylitol productivity, and investigate whether the industrial background inflicts any difference in the xylose metabolism. Specifically, we evaluate the hypothesis that aldehyde-resistant SA-1 has a better redox environment, since most reductases that take part in HMF or furfural detoxification reactions are NADPH-dependent - the same cofactor required by XR activity -, and therefore represent a good chassis for metabolic engineering envisioning the production of reduced sugars.

Before proceeding to an in-depth analysis of xylitol productivity in the transformed strains, a preliminary assay to assess CENPKX, JAYX, and FMYX performance on xylose-containing media was carried out (**Figure 3**). Oxygen-limiting batch cultivation was performed in order to mimic the cultivation conditions typically performed in the Brazilian E2G industry [36], aiming at the possibility of using the same operational infrastructure for xylitol production. After 120h of semi-anaerobic cultivation on YPDX, CENPKX and FMYX produced similar concentrations of xylitol (6.08 g/L and 5.57 g/L, respectively), while JAYX had limited production of 2.60 g/L (**Figure 3A**). Even though all strains showed substantially superior xylitol productivity in relation to their wild-type counterparts, PE-2 performance was surprisingly the lowest. Next, strains CENPKX and FMYX were evaluated for xylitol production in the presence of (HMF (**Figure 3B**). FMYX was able to maintain productivity even when 2 g/L of the aldehyde is present in the medium, while the laboratory failed to keep up with the xylitol titre obtained in control. This result, together with the low xylitol production observed in PE-2, were key to choosing FMYX for the next experiments.

**Figure 3:**
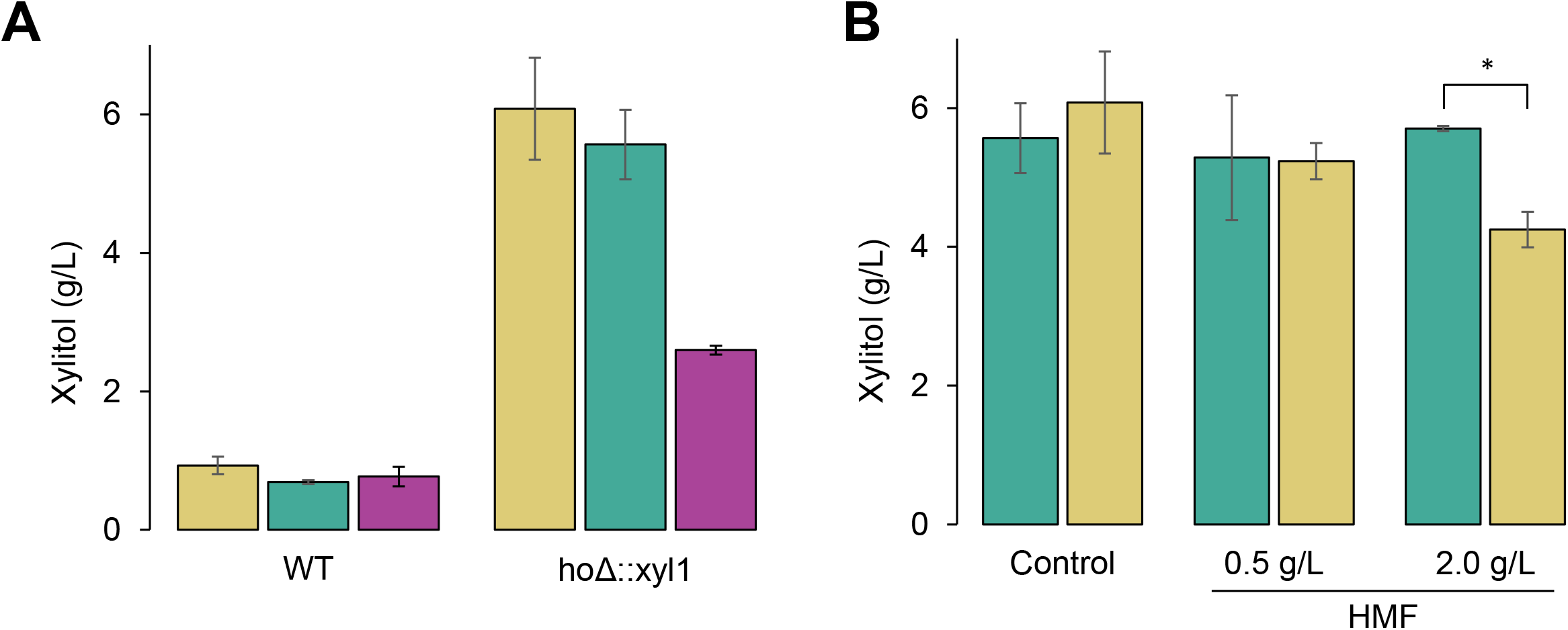
A prospect of xylitol productivity in engineered industrial *S. cerevisiae* strains. **A.** Xylitol productivity of wild-type and xylose reductase-expressing (*ho*Δ::xyl1) strains PE-2, SA-1 and laboratory CEN.PK-122 in semi-anaerobic batch fermentation with YPDX (0.5/2%) after 120 hours. **B.** Xylitol productivity in the presence of varying HMF concentration of strains FMYX and CENPKX in semi-anaerobic batch fermentation with YPDX (0.5/2%) after 120 hours. Asterisk (*) represents statistically different averages between bars (p-value < 0.05).

### Industrial background favors xylitol production in *XYL1* expressing yeast

Here we explore how the xylose:glucose ratio, aeration conditions and cell inoculum can affect xylitol production of strains FMYX and CENPKX in batch cultivation. Such evaluations have been traditionally documented in genetically modified *S. cerevisiae* [37], and re-analyzed in this study in order to understand if the strains’ backgrounds influence xylitol productivity - given that the same genomic edition was performed in both. Xylose consumption and xylitol production graphs are presented in **Figure 4** and the complete dataset in **Table 2**.

**Figure 4:**
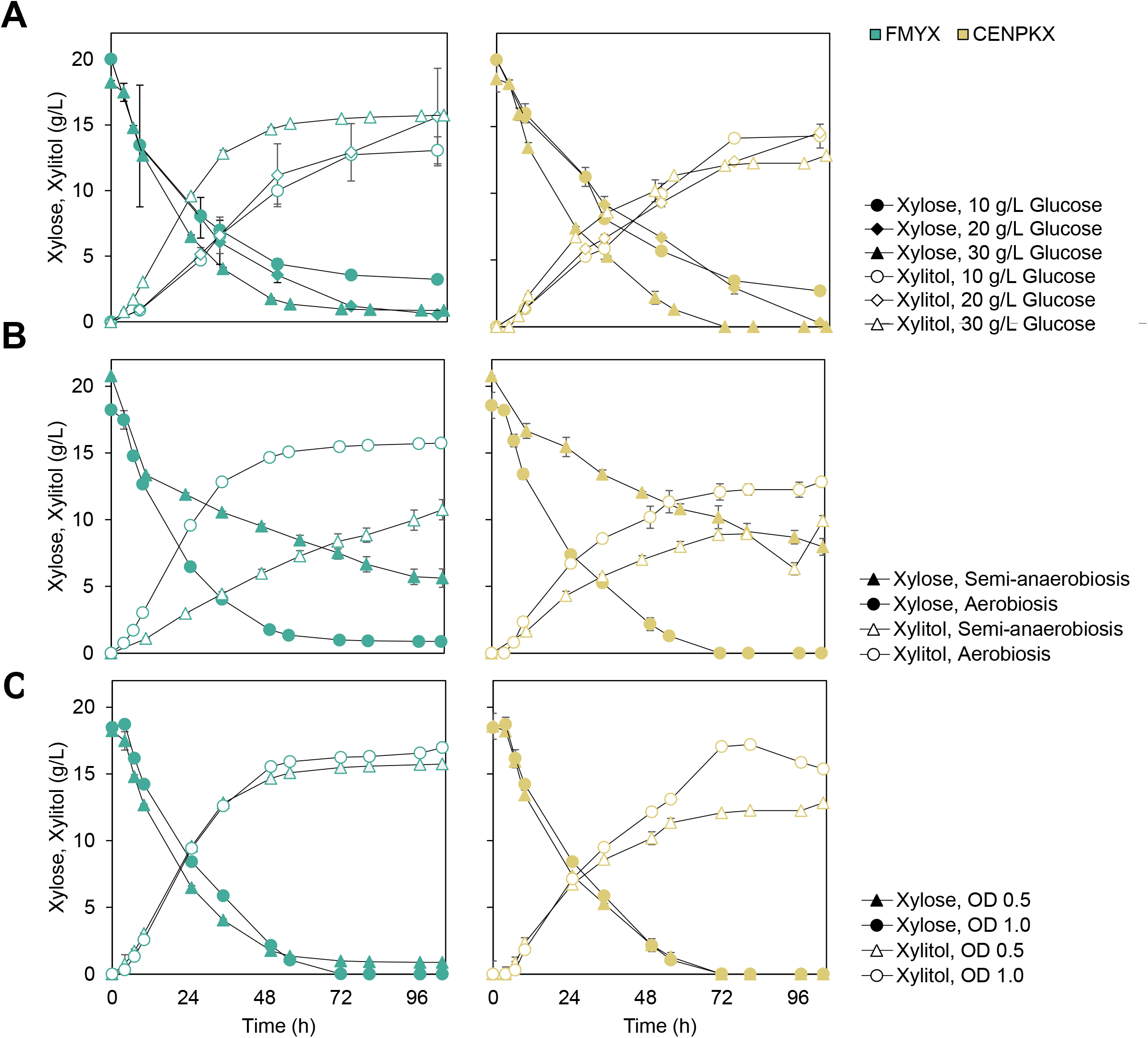
Xylitol productivity of strains FMYX (green on the left) and CENPKX (yellow on the right) in varying cultivation conditions. Filled symbols (•/□/▲) represent xylose and empty (◦/□/□), xylitol. **A.** Aerobic cultivation with an initial optical density (OD_600nm_) of 0.5, 20 g/L xylose and different glucose concentrations: 10 g/L (•/◦), 20 g/L(□/□) or 30 g/L (▲/□) of the cosubstrate. **B.** Cultivation in 20 g/L xylose, 30 g/L glucose, initial OD_600nm_ of 0.5 and varying oxygenation scenarios: semi-anaerobic (▲/□) and aerobic (•/◦). **C.** Aerobic cultivation with 20 g/L xylose, 30 g/L glucose and different cell mass for inoculum: OD_600nm_ 0.5 (▲/□) or OD_600nm_ 1.0 (•/◦).

**Table 2:**
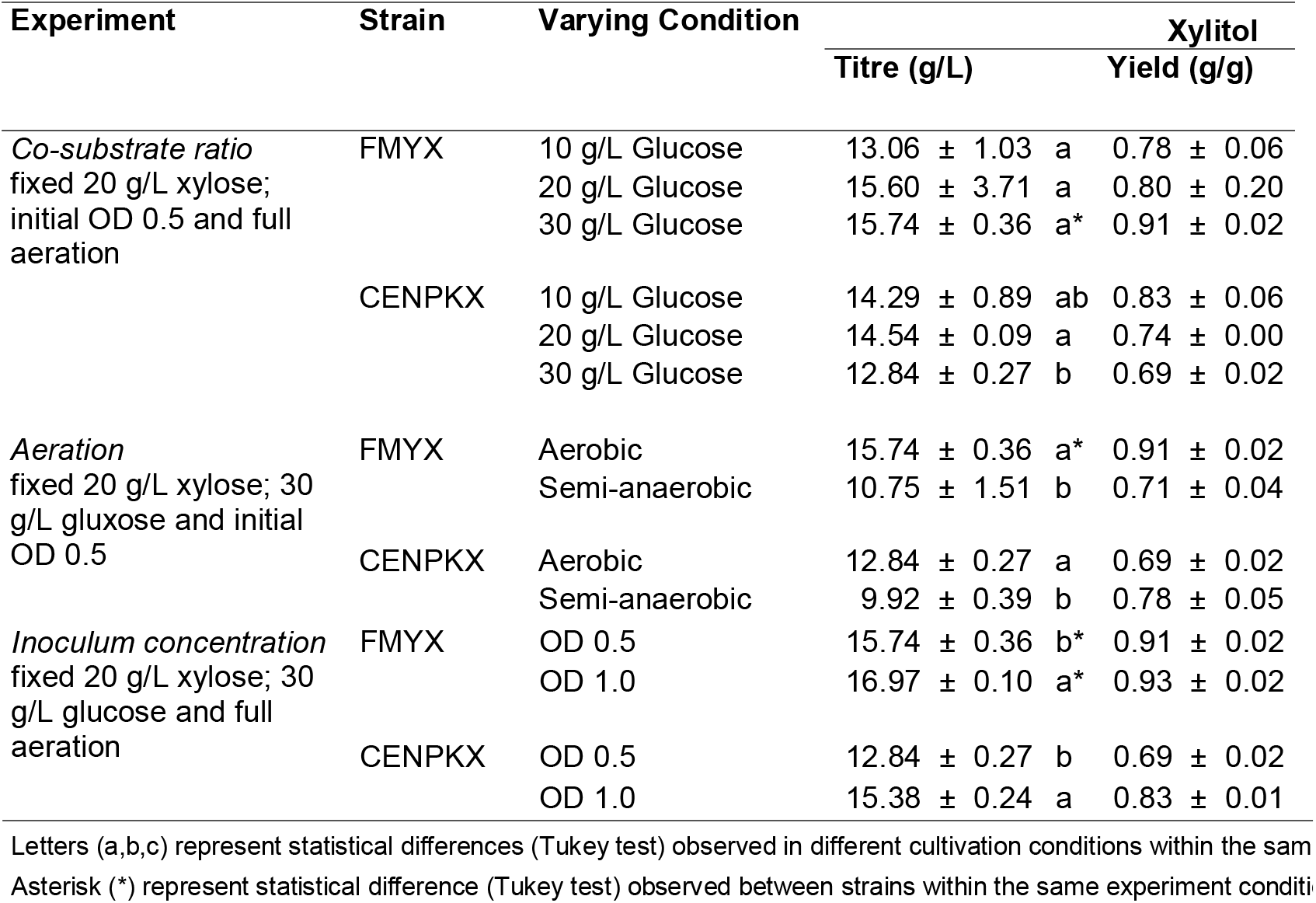
Xylitol productivity of strains FMYX and CENPKX in YPDX under varying cultivation con

First, to understand the effect of co-substrate availability in xylitol production in modified SA-1 and CEN.PK-122, the strains were submitted to cultivation assays with different xylose:glucose ratios. The assay was carried out with 20g/L xylose, full aeration, an initial OD_600nm_ of 0.5 and varying concentrations of glucose (10, 20 and 30 g/L) (**Figure 4A**). For both strains, 10 g/L of glucose prevented the complete consumption of xylose, and a residual amount of the last is present after 104 hours of cultivation. Higher concentrations of the co-substrate allowed the complete metabolization of xylose by both strains. Regarding xylitol production, laboratory strain CENPKX peaks at 14.54 ± 0.09 g/L for the cultivation with 20 g/L glucose, while FMYX has the best overall performance with 30 g/L of the co-substrate, producing 15.74 ± 0.36 g/L xylitol.

Following, we tested xylitol productivity performance of transformed strains in semi-anaerobic cultivations (**Figure 4B**), maintaining an initial OD_600nm_ of 0.5 and a ratio of 20:30 (xylose:glucose) concentration. Higher glucose concentration was chosen to resemble typical sugarcane hydrolysate. The oxygen-limiting environment resulted in lower xylitol titre (10.75 ± 0.75 g/L) for FMYX in comparison to the aerated cultivation and statistically different xylitol yields (0.91 and 0.71, respectively). At the same time, CENPKX reduced its overall xylitol titre, but no difference was observed in this sugar yield between the different oxygenation cultivation scenarios.

Finally, to test whether the initial inoculum affects xylitol productivity of strains CENPKX and FMYX, we have cultivated both strains with 30 g/L glucose, 20 g/L xylose, full aeration, and varied initial OD_600nm_ - 0.5 or 1.0 (**Figure 4C**). For both strains, we observed the same behavior: higher initial cell concentration allowed higher xylitol titre. For CENPKX, xylitol concentration after 104 h cultivation increased from 12.83 ± 0.27 g/L (OD_600nm_ 0.5) to 15.38 ± 0.24 g/L (OD_600nm_ 1.0). Meanwhile, industrial FMYX was able to produce 15.74 ± 0.36 g/L and 16.97 ± 0.10 g/L xylitol when OD_600nm_ was 0.5 and 1.0, respectively. Again, FMYX outperformed CENPKX in both scenarios.

### Batch fermentation of raw sugarcane bagasse hydrolysate for xylitol production

We then proceeded to evaluate the possibility of using sugarcane bagasse hydrolysate - traditionally used in the Brazilian E2G industry [36] - to produce xylitol using strains FMYX and CENPK.

Sugarcane straw was donated by GranBio SA (Bioflex plant) and hydrolysis was performed with Cellic CTec3. Cells were further inoculated at OD_600nm_ 1 and cultivated with aeration. Xylitol productivity was assessed in the same operational process conditions as is in E2G: batch fermentation of non-detoxified hydrolysate with remaining solid phase [36] (**Figure 5**). The hydrolysate composition is presented in **Table 3** and xylitol productivity of FMYX and CENPKX, in **Table 4**. The harsh fermentation conditions revealed a challenge for xylitol productivity in both strains, yielding 3.65 ± 0.16 g/L and 1.04 ± 0.45 g/L (p-value = 7.0E-04) for FMYX and CENPK, respectively. In fact, CENPKX was not even able to consume glucose after 120h of fermentation, while FMYX did metabolize all the co-substrate and some of the xylose available.

**Table 3:** Sugarcane bagasse hydrolysate composition.

| Analyte | Composition (g/L) |
| --- | --- |
| Glucose | 49.87 ± 7.12 |
| Xylose | 23.95 ± 3.66 |
| Acetic Acid | 5.84 ± 0.78 |
| Formic Acid | 1.33 ± 0.27 |
| HMF | 0.59 ± 0.01 |
| Furfural | 1.20 ± 0.07 |

**Table 4:**
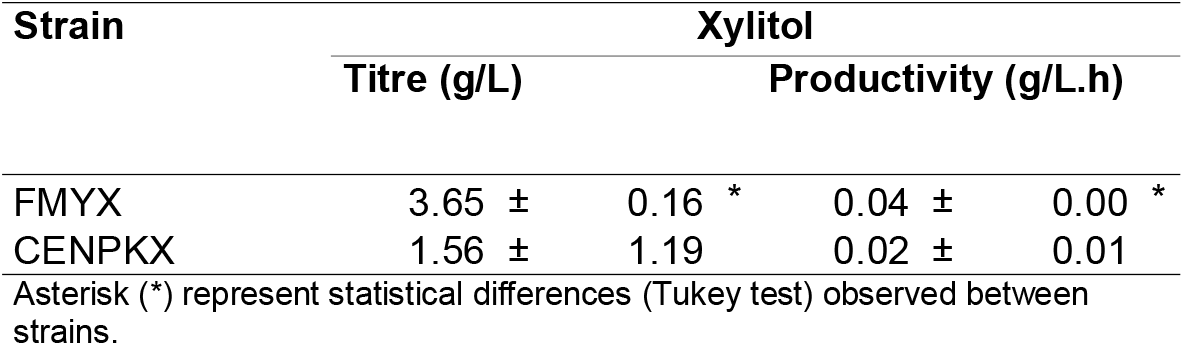
Xylitol productivity of strains FMYX and CENPKX in sugarcane bagasse hydrolysate.

**Figure 5:**
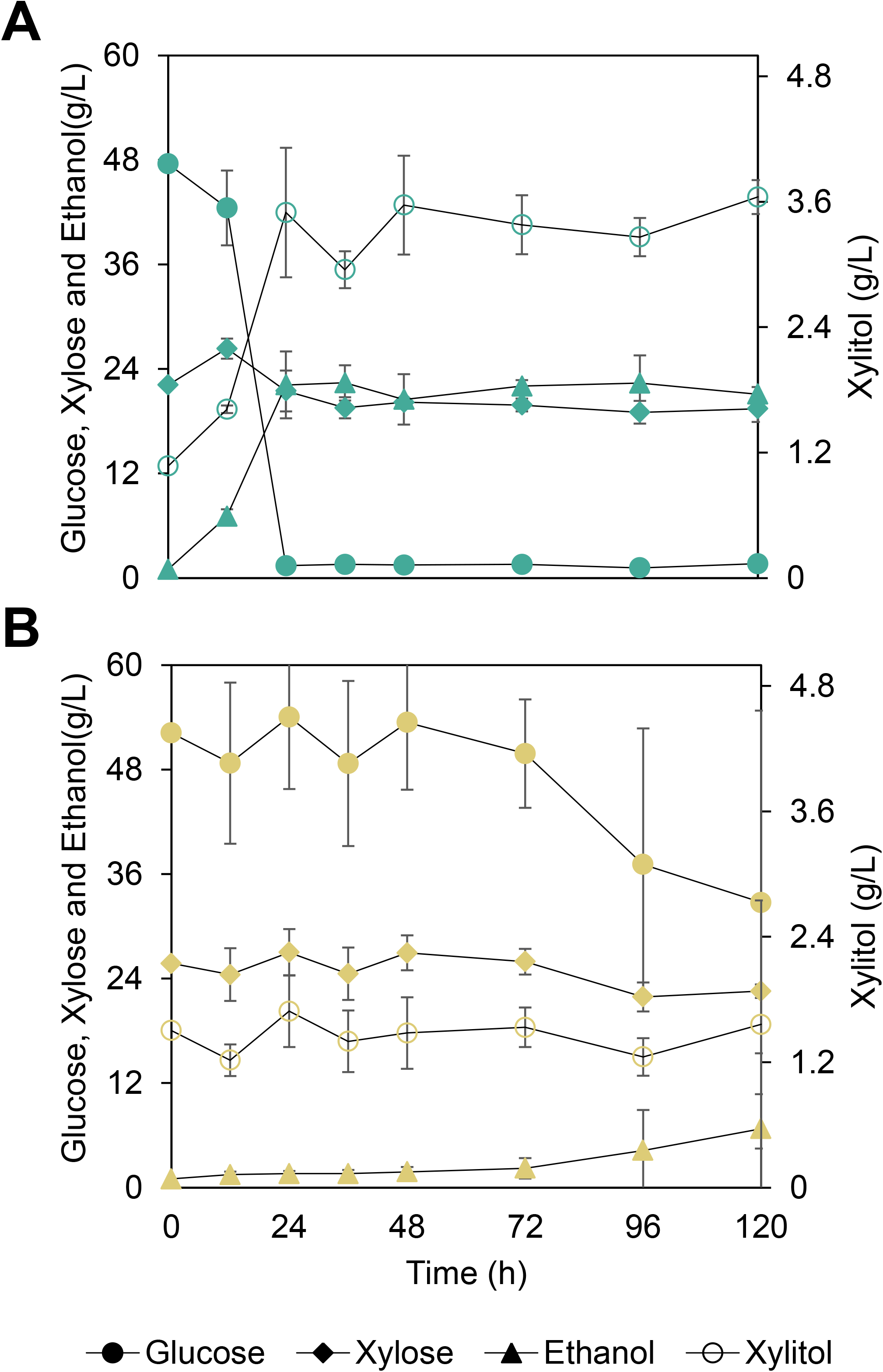
Batch fermentation of raw sugarcane bagasse hydrolysate with remaining solid particles and full aeration for xylitol production by strains FMYX (**A**) and CENPKX (**B**). Curves represent consumption of glucose (•) and xylose (□), or production of ethanol (▲) and xylitol (◦).

### The redox environment of SA-1 and CEN.PK-122

Next, we sought to uncover why strain FMYX outperformed CENPKX regarding xylitol production, especially in stress-free scenarios. Because FMYX and CENPKX have just one copy of *XYL1* integrated in the genome, higher xylitol titres obtained by the first must be related to the strains’ background prior to the genomic editing. Complete metabolization of xylose could explain the diverging results. Although *S. cerevisiae* is known to have genes homologous to xylose dehydrogenase (XDH) enzymes (such as *XYL2*), there is conflicting evidence whether xylose can induce its activity in converting xylitol to xylulose [38]. Even though xylose conversion to ethanol could have impaired CENPKX’s xylitol productivity, maximum alcohol titre produced by the strain is 11.53 ± 0.35 g/L, compared to 12.66 ± 0.11 g/L by FMYX, in the last cultivation condition tested. Therefore, this fails to explain the performance difference between strains.

As previously stated, cofactor regeneration is crucial for xylitol production in *S. cerevisiae*, given that commonly used reductases are NADPH-dependent and xylose metabolism is incomplete for that goal. Accordingly, we moved on to test whether there is a contrast in reduced cofactors that could enhance XR activity and further xylose conversion in FMYX. Relative NADPH in exponential growth at optimal conditions was assessed by a luminescence assay using the transformed and wild-type strains - SA-1 and CEN.PK-122 (**Figure 6**). The results confirm that industrial SA-1 has 33% more NADPH than laboratory CEN.PK-122 (p-value = 3.0E-04), and that *xyl1* genomic integration has not affected cofactor availability in FMYX while CENPKX fell behind in performance compared to its wild-type counterpart. CENPKX has only 59% of the amount of cofactor available in CEN.PK-122 (p-value < 1.0E-04).

**Figure 6:**
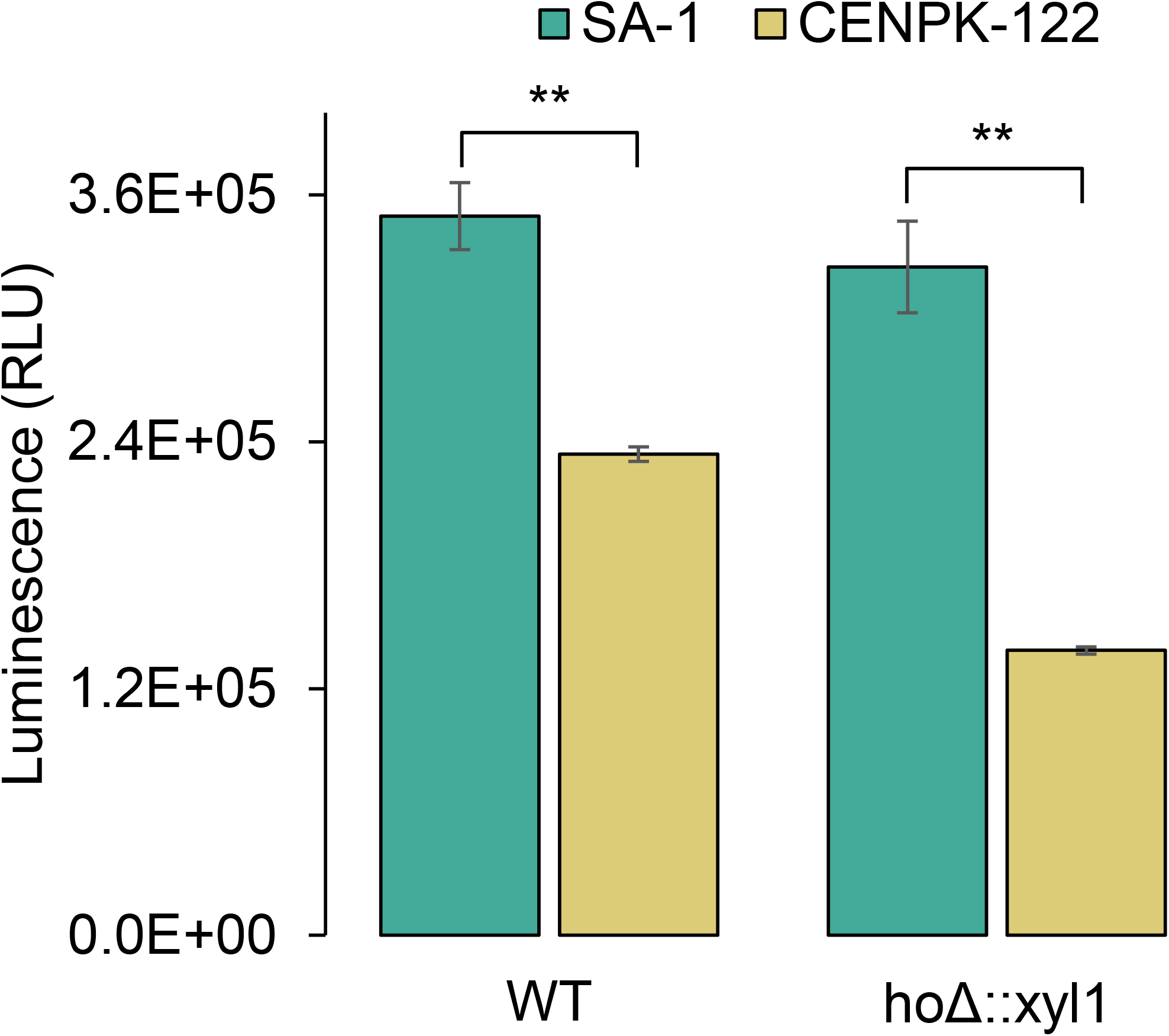
Relative NADPH quantification using a luminescence assay for wild-type (WT) and genetically modified (*ho*Δ::xyl1) strains SA-1 and CEN.PK-122. Asterisks (**) represent statistically different averages between bars (p-value < 0.001).

## 1. DISCUSSION

In summary, our results show the development of a functional CRISPR engineering method for diploid strains PE-2 and SA-1 and set the basis for xylitol production using sugarcane bagasse, while unraveling the rationale behind the excellent performance of the industrial FMYX. Engineering such widely used bioethanol yeast is relevant given that their robust background and adaptability to harsh fermentation conditions result in recombinant S. cerevisiae suitable for industrial applications. In this context, commercial bio-fabrication of xylitol using S. cerevisiae stands out as a promising alternative to the chemical route.

When it comes to CRISPR-engineering of the Brazilian bioethanol strains, the developed sgRNA prospection confirms the noteworthy genetic difference between tailored industrial strains and the S288c model, backing up the necessity of analyzing sgRNA efficiency in different yeast strains in addition to using the available prediction tools. For both strains analyzed here, a total of 14% of the sequences predicted to be efficient in the S288c model had high chances of leading to off-target Cas9 activity. Even though high accounts of genetic diversity and complex population structure in the so-called budding yeast is notorious for the scientific community [39], the use of the S288c model as the sole source of genetic information for many important inquiries and assays remains true. When it comes to sgRNA prospection, coupling the available sgRNA-prediction software with in-depth investigations into relevant strains’ genome comes in hand to guarantee efficient rational editing.

Regarding the assembly and transformation of the editing-parts, improvements allowed for fast and cheap protocols with high editing rates. Even though many single plasmid based CRISPR strategies have been developed for S. cerevisiae [40-45], they differ greatly when it comes to sgRNA insertion strategies and overall editing efficiency. In our system, the single-restriction methodology of the pGS plasmid allows for the fast exchange of sgRNAs. Additionally, when applied in co-transformation with a donor DNA, sgRNA-containing pGS induced high percentages of editing effectiveness - above 97% when 1500 ng of the vector and 1000 ng of the donor DNA were applied.

While high concentration of donor DNA is crucial for the general efficiency of the editing system, pGS concentration plays a decisive role in the final count of edited colonies. The highest donor DNAs concentration tested (1 μg, 16.83pmols) led to efficiency rates very close to 100% in disrespect of the plasmid concentration, while low concentrations often led to low-efficiency rates. On the other hand, transformation events bearing the highest plasmid concentration (1.5 μg, 0.23 pmols) led to a generally greater number of edited colonies, when comparing across the same donor concentrations. This is especially true for co-transformation with 1 and 0.5 μg of donor DNA. Interestingly, moderate concentrations of the sgRNA and Cas9-bearing plasmid (100ng-500ng), which are commonly applied in S. cerevisisae genome editing, were found not to be as effective in our strains. These results are true for the wild-type diploid PE-2 as well as a haploid derivative of the strain. Although polyploid strains are more suitable for industrial fermentation, their segregants are typically used for the construction of robust hybrids [46,47], herein the importance of the system effectiveness in haploid strains of industrial relevance.

After establishing optimal conditions for genome editing with the CRISPR-Cas9 system, PE-2 and SA-1 were put to test. High titre xylitol production in indigenous yeast commonly requires the expression of heterologous enzymes, since the endogenous aldose reductase (coded by gene GRE3) displays low rates of d-xylose catabolism. Although there are reports of chromosomal integration of xylose reductase in *S. cerevisiae*, no work describes the use of a CRISPR-Cas9 system for this purpose. Also, genome editing of the relevant industrial strain SA-1 is first described in this study.

The HO locus – responsible for the mating-type switch in S. cerevisiae – was chosen for the introduction of XYL1. Prior to the insertion, HO’s documented inert function in cell fitness [48] was confirmed for the bioethanol strains used in this work. A SNP in HO between the industrial yeasts and the laboratory CEN.PK-122 demanded specific sgRNA sequences for successful Cas9 activity in each strain. This finding corroborates with the importance of sgRNA screening in non-conventional *S. cerevisiae* prior to their genetic engineering. Again, sgRNAs suggested by popular design tools based on model yeast genome would fail to be functional, leading to time-consuming sgRNA survey. The database provided in this study shortens the efforts for sgRNA efficiency-check in major strains PE-2 and SA-1.

Prospect of xylitol productivity in mutant strains in oxygen-limiting batch fermentation - scenario typically found in the Brazilian E2G industry [36] - revealed that PE-2’s performance was restricted compared to the other strains. Previous report on the use of a GRE3 episomal overexpressing PE-2 in xylitol production with aeration [15] shows good performance of this chassis, suggesting that the semi-anaerobic environment hampered JAYX execution. Testing of xylitol productivity in presence of HMF revealed that edited SA-1 was able to maintain its performance when 2 g/L of the aldehyde was present in the medium, while CENPKX failed to keep up with the xylitol titre obtained in the control condition. 2 g/L HMF have been described as highly damaging to xylitol production in oxygen-limiting xylitol batch fermentation [49], confirming the outstanding FMYX performance. Besides assessing productivity in conditions with fermentation inhibitors that resemble lignocellulosic hydrolysates, xylitol production in the presence of HMF also corroborates with the hypothesis that SA-1 has a better redox environment. Bearing in mind that formyl detoxification is conducted by multiple NADPH-dependent aldehyde reductases (e.g., ARI1, ADH6, ADH7) [50], HMF-resistant strains might be suitable for better XR activity and, consequently, xylitol production.

The influence of varying co-substrate ratio, aeration and initial cell density on xylitol productivity have already been examined in other strains [18,37], and here revisited to explore the different chassis effect over xylose metabolization. Varying cultivation parameters in optimal conditions were tested to evaluate how the genetic background influences xylitol productivity in strains CENPKX and FMYX. Because the sole expression of genes encoding XR prevents the use of xylose as a carbon source for energy maintenance, the regeneration of cofactors and cell growth is dependent on co-substrates - such as glucose. At the same time, xylitol productivity in mutant *S. cerevisiae* relies on the availability of NADPH - regenerated in the oxidative part of pentose phosphate pathway (PPP) - for the activity of a reductase.

While xylose uptake into the cells is inhibited by glucose [51], we chose this co-substrate to mimic the conditions in sugarcane hydrolysate fermentation, where this is the most prominent carbon source for *S. cerevisiae*. A fixed concentration of 20 g/L xylose was used throughout this assay to achieve a higher turnover rate for the reductase and guarantee a higher xylitol formation rate [37]. When altering glucose concentration regarding xylitol, industrial FMYX presented a better overall xylitol productivity in the same conditions evaluated for the laboratory. CENPKX underperforms when 30 g/L of the co-substrate is present in the medium, indicating a possible glucose repression scenario. Indeed, 20:30 xylose:glucose ratio is the closest to an industrial scale fermentation [52], substantiating that FMYX should be more appropriate for this purpose.

Regarding oxygen availability, Hallborn *et. al* [37] have previously suggested that limited oxygen supply favors xylitol production because less NAD(P)H is spent on ATP production. Nevertheless, our results show the opposite - especially for industrial FMYX -, proposing that the yeast background indeed plays an important role in xylitol productivity. In fact, further work on xylitol production by recombinant *S. cerevisiae* usually apply aeration for xylose metabolization [13], supporting our findings. The difference of xylitol titre between strains in semi-anaerobiosis again reveals a tendency of industrial FMYX to produce more of the reduced sugar.

The positive effect of higher initial cell density in xylitol productivity has been previously reported by Kogje and Ghosalkar (2017) [18], corroborating with our results. More yeast cells guarantee that the co-substrate is fully metabolized through the PPP and NADPH is regenerated for the xylose reductase activity. Overall, in the conditions tested, edited industrial SA-1 outperformed the laboratory CEN.PK-122 expressing a XR, confirming that the genetic background plays a key role in xylitol productivity, even in a stress-free cultivation environment.

Although bio-based xylitol production has been widely explored, sugarcane bagasse fermentation using *S. cerevisiae* has not yet been documented for this purpose. One of the challenges of using sugarcane hydrolysate in whole-cell fermentation resides in the inhibitory nature of several pretreatment by-products. While such hemicellulosic feedstock has been tested for xylitol production using yeast *Candida guilliermondii* [53,54], approaches using recombinant *S. cerevisiae* have been limited to either corn cob [15,17,18,55], rice straw or woody biomass [16,56] as natural xylose sources. Using sugarcane biomass to produce high added-value chemicals with *S. cerevisiae* is a prospect for future biorefineries [57], and our experiments join this discussion.

For xylitol production using sugarcane bagasse, similar conditions applied in the Brazilian industrial E2G were used in order to simulate the performance of the existing infrastructure. Batch fermentation of raw non-detoxified acid-pretreated bagasse hydrolysate with remaining solid particles revealed challenges for xylitol productivity of both FMYX and CENPKX, still the first was able to produce three times more xylitol than the laboratory. These results dialogue with what we have observed in optimal conditions fermentation, regarding glucose repression in CENPKX and better overall performance of FMYX. Here, besides the intrinsic ability of FMYX to produce more xylitol compared to CENPKX, the documented robustness of the industrial strain towards second-generation inhibitors also contributes to its achievement. Nevertheless, the recalcitrant hydrolysate with remaining solid particles prevented the full metabolization of xylose by FMYX, and operational improvements are presumed necessary.

Finally, an investigation to understand the contrasting performance regarding xylitol productivity between FMYX and CENPKX was performed. While superior xylitol titre produced by FMYX in the sugarcane bagasse hydrolysate might be related to the strain’s robustness towards fermentation inhibitors [22], this shouldn’t elucidate why the industrial background outperformed CENPKX in optimal cultivation conditions. Xylitol conversion to xylulose through an endogenous XDH activity could reduce xylitol yields, once xylose would be further metabolized into ethanol. Even so, higher concentrations of this alcohol were produced by FMYX, suggesting that *XYL2* activity did not impair CENPKX’s xylitol production. At last, quantification of cofactor NADPH - participating in XR activity - revealed an outstanding higher availability of this cofactor in the industrial FMYX.

While efforts have been made to engineer cofactor preference in XR [15] or even to increase the flux through PPP by overexpressing a glucose-6-phosphate dehydrogenase to produce more NADPH [58], we deduce that tailored yeast strains might have an endogenous favorable redox potential. Therefore, industrial SA-1 represents a good chassis for genetic engineering not only for the already known robust phenotype, but also for the possibility of achieving better yields of reduced sugars catabolized by NADPH-dependent reductases.

## CONCLUSION

This work has enabled the development and standardization of an efficient CRISPR-Cas9-based method for the metabolic engineering of industrial diploid *S. cerevisiae* strains applied in the bioethanol industry. Apart from the proof of concept of the method developed during this work - based on the *URA3* gene deletion and insertion into the PE-2 strain - this approach has been successfully used for efficient xylitol production in strains PE-2, SA-1, and CEN.PK-122. We have observed that the industrial background enabled a better xylitol productivity, in comparison to the laboratory control. Growth of transformed SA-1 and CEN.PK-122 in YPDX revealed that in cofactor-limiting scenarios, the first outperformed the second. NADPH quantification indicated a superior redox environment in SA-1, suggesting that yeast strains applied in harsh industrial processes might have a better cofactor regeneration trait and are good chassis for metabolic engineering. Finally, it is important to note that fermentation of raw sugarcane bagasse hydrolysate did not result in good xylitol yields, indicating a technological bottleneck for the use of *S. cerevisiae* in the same operational conditions that E2G is now produced.

## LIST OF ABBREVIATIONS

CRISPR: Clustered Regularly Interspaced Short Palindromic Repeats
Cas9: CRISPR associated protein 9
dsOligos: donor DNAs
E2G: Second-Generation Bioethanol
HMF: 5-hydroxymethylfurfural
HPLC: High performance liquid chromatography
LB: Luria-Bertani
KO: Knock-out
NoNAG: NGG PAM sequences
nt: nucleotide
OD: Optical density
PCR: Polymerase chain reactions
PE-2: Pedra-2
sgRNA: Single-guided RNA
SNP: Single nucleotide polymorphisms
SS: Salmon Sperm
YPD: Yeast Extract Peptone and Dextrose
XR: Xylose Reductase

## AUTHOR’S CONTRIBUTION

GT conceived the present idea and FM conceptualized the paper. GT and GP supervised the study. FM, CM, AC and GT designed the experimental outline. CM and SN worked on the sgRNA analysis. CM carried out the CRISPR protocol procedures. CM, FM and FS produced transformed strains. FM, FS and BV performed the xylose cultivation in defined media. FM performed the NADPH quantification assay. CV, TS and FM did the sugarcane hydrolysate fermentation. FS, CV, TS, MS and RG helped with HPLC analysis. FM and CM wrote the manuscript, with inputs from CV. All authors read and approved the final manuscript.

## DECLARATION OF COMPETING INTEREST

The authors have no conflicts of interest to declare.

## AVAILABILITY OF DATA AND MATERIALS

The datasets used and/or analyzed during the current study are available from the corresponding author on reasonable request.

## ACKNOWLEDGMENTS

We want to thank Prof. Dr. Marcelo Carazzolle from The University of Campinas (Brazil) for assisting the sgRNA analysis in PE-2 and SA-1 genomes.

## FUNDING

This study was financed by the *Fundação de Amparo à Pesquisa do Estado de São Paulo* (FAPESP, São Paulo, Brazil) through a scholarship to FM (grant number 2015/06677-8), CM (grant number 2018/03403–2) and GS (BIOEN grant number 2016/02506-7).

## ETHICS APPROVAL AND CONSENT TO PARTICIPATE

Not applicable.

## CONSENT FOR PUBLICATION

Not applicable.

## ADDITIONAL FILES

**Additional File 1**

**Format:** Table (XLS)

**Title: Supplementary Table S1**

**Description:** Main primers used in this study.

**Additional File 2**

**Format:** Table (CSV)

**Title: Supplementary Table S2**

**Description:** sgRNA prospect in PE-2.

**Additional File 3**

**Format:** Table (CSV)

**Title: Supplementary Table S3**

**Description:** sgRNA prospect in SA-1.

**Additional File 4**

**Format:** Image (PDF)

**Title: Supplementary Figure S1**

**Description:** Cloning procedures for using pGS in a CRISPR-Cas9 editing event.

**Additional File 5**

**Format:** Image (PDF)

**Title: Supplementary Figure S2**

**Description:** Transformation efficiency of the *LEU2 locus* knockout in LVY34.4 (PE-2, MATa).

**Additional File 6**

**Format:** Image (PDF)

**Title: Supplementary Figure S3**

**Description:** Testing of sgRNAs targeting the *HO locus* in strains CEN.PK-122, SA-1 and PE-2.

**Additional File 7**

**Format:** Image (PDF)

**Title: Supplementary Figure S4**

**Description:** Testing of the fitness of strains PE-2, SA-1 and CEN.PK-122 with a *ho*Δ genotype.

**Additional File 8**

**Format:** Image (PDF)

**Title: Supplementary Figure S5**

**Description:** Editing efficiency of strains PE-2, CEN.PK-122 and FMY097/FMY034 (SA-1 segregants) for integration of a xylose reductase cassette.

## Notes

### Competing Interest Statement

The authors have declared no competing interest.

