## Supplementary figures and images for "Rational engineering of industrial *S. cerevisiae*: towards xylitol production from sugarcane bagasse"

### Figure S1

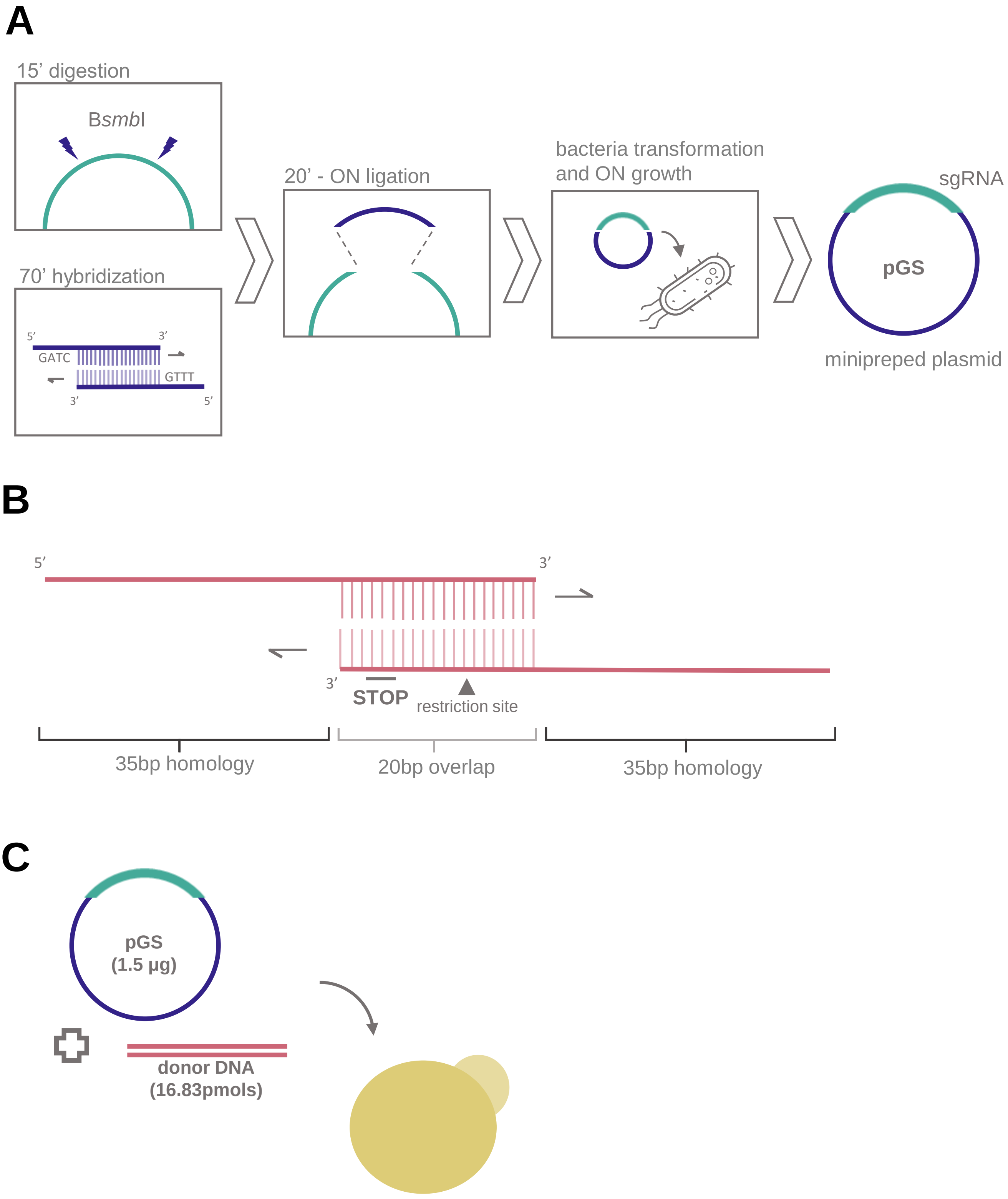

### Figure S2

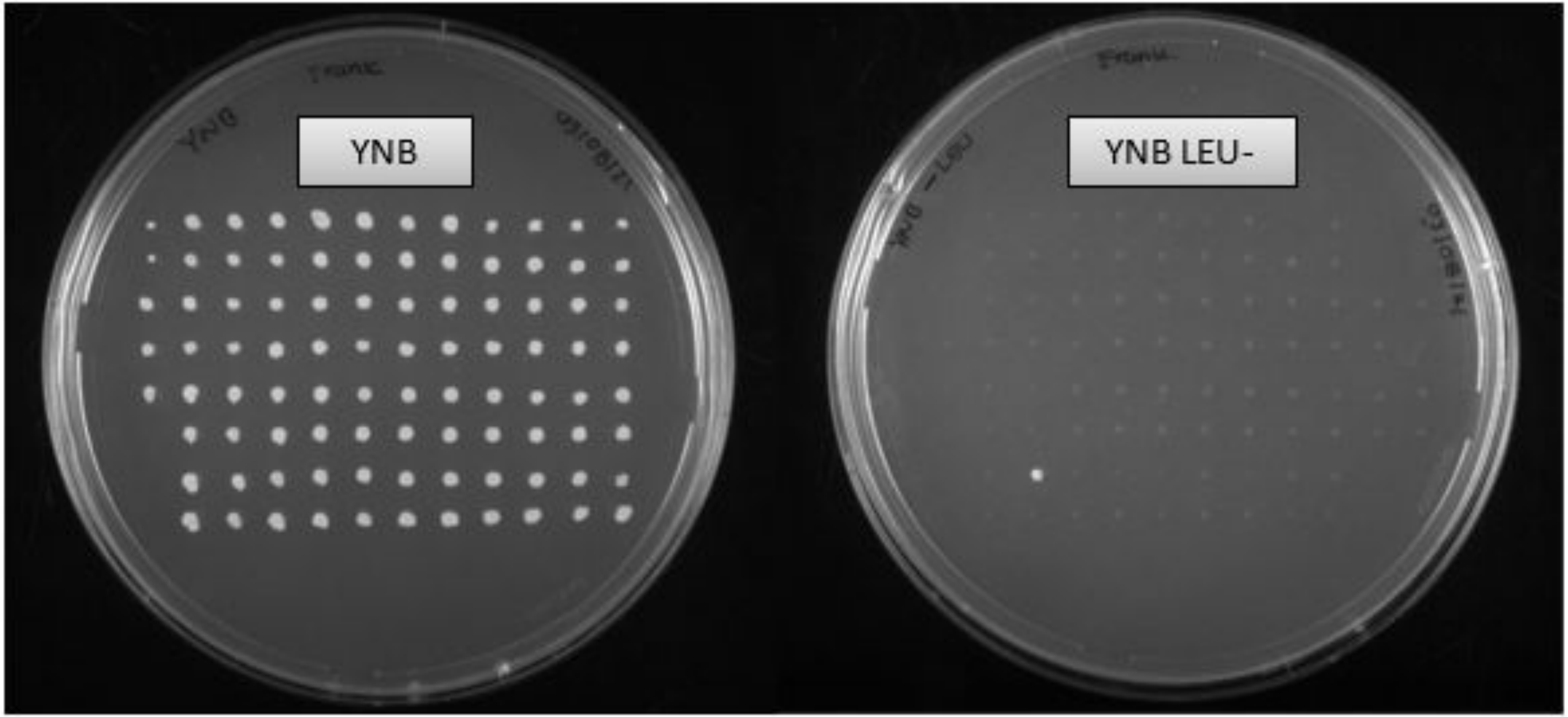

### Figure S3

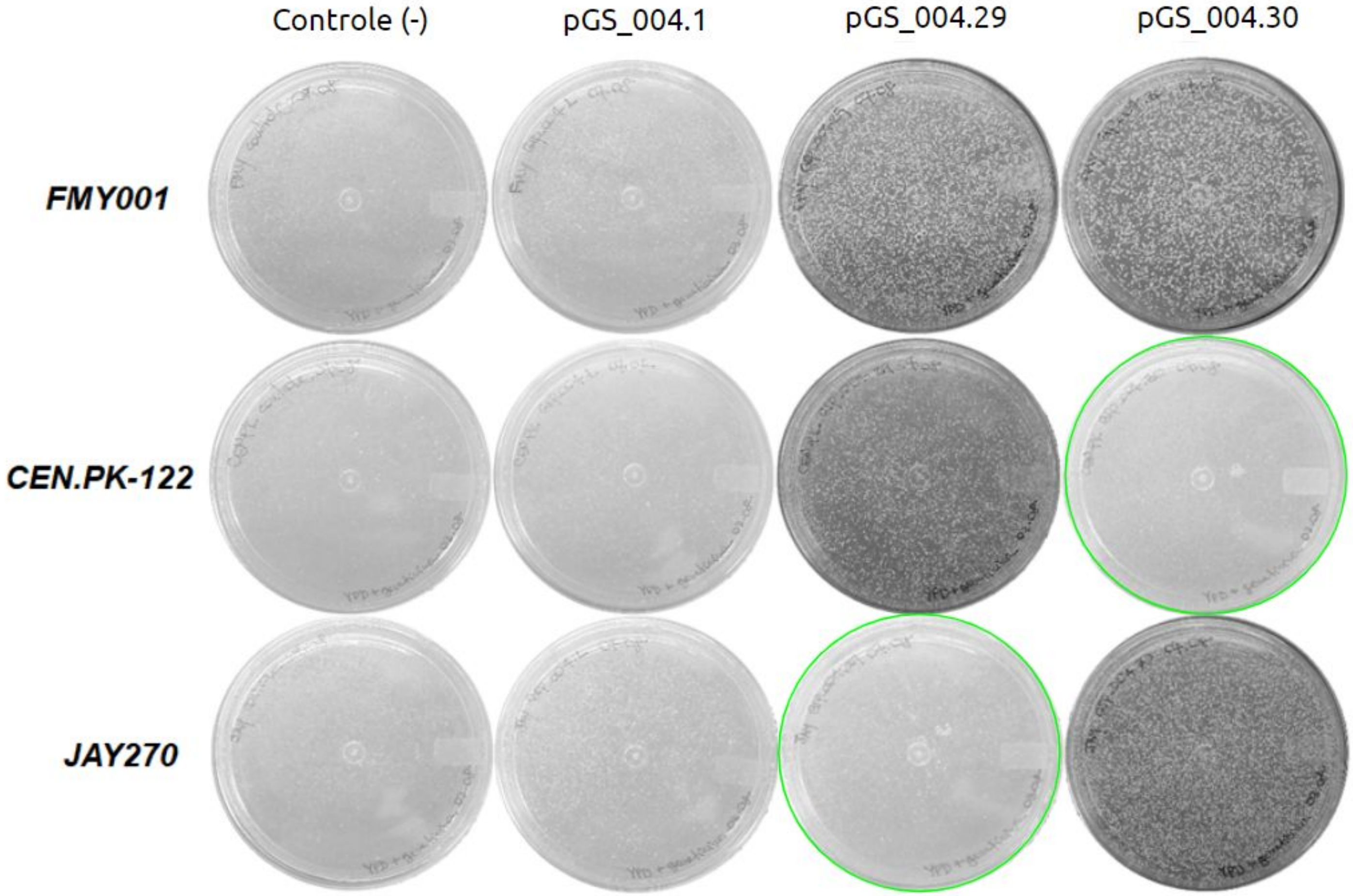

### Figure S4

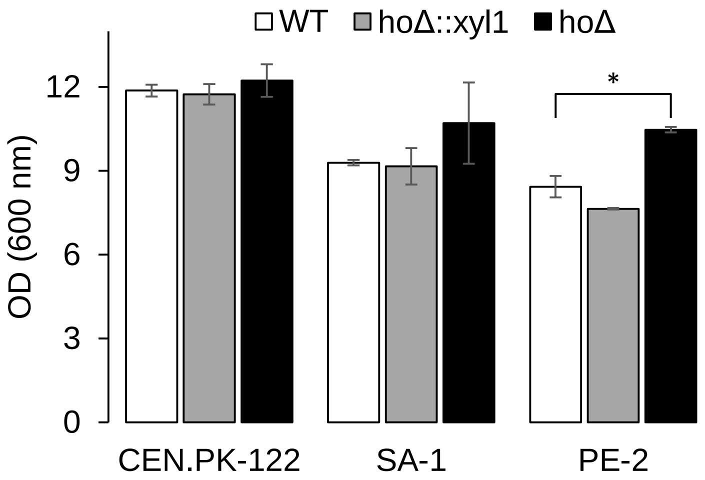

### Figure S5

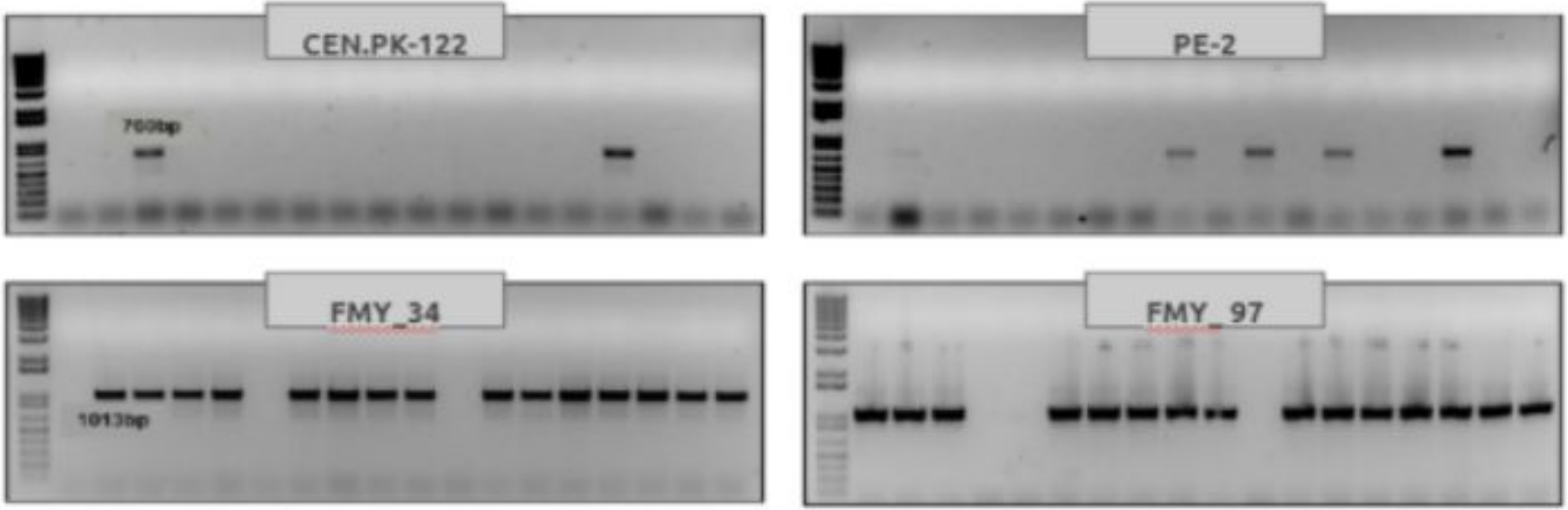
