## Supplementary Table S1 for "Rational engineering of industrial *S. cerevisiae*: towards xylitol production from sugarcane bagasse"

| Primers | Sequence (5’->3’) | Purpose |
|---|---|---|
| SgRNAseq_F | CGTCTTCTCTTTGAAAAGATA | Sequencing of sgRNAs in pGS |
| SgRNAseq_R | TTGAAGTCCGTTTATTAACTT | Sequencing of sgRNAs in pGS |
| 01_F | GATCGCACACGGTGTGGTGGGCCC | Hybridization of a dsOligo contaning a sgRNA sequence targeting the URA3 gene (KO) |
| 01_R | AAACGGGCCCACCACACCGTGTGC | Hybridization of a dsOligo contaning a sgRNA sequence targeting the URA3 gene (KO) |
| URA3KO_F | CAGAATAGCAGAATGGGCAGACATTACGAATGC ACACGGTGTGGTGGGCTGATCA | Hybridization of the donor DNA for the URA3 gene KO |
| URA3KO_R | CTTCTTCCGCCGCCTGCTTCAAACCGCTAACAA TATGATCAGCCCACCACACCGT | Hybridization of the donor DNA for the URA3 gene KO |
| 02_F | GATCGAAGTAACAAAGGAACCTAG | Hybridization of a dsOligo contaning a sgRNA sequence targeting the URA3 gene (KI) |
| 02_R | AAACCTAGGTTCCTTTGTTACTTC | Hybridization of a dsOligo contaning a sgRNA sequence targeting the URA3 gene (KI) |
| URA3KI60_F | TGCCCAGTATTCTTAACCCA | Amplification of the URA3 gene plus 60bp homologies |
| URA3KI60_R | TTAAATTGAAGCTCTAATTTGTGA | Amplification of the URA3 gene plus 60bp homologies |
| URA3KI1000_F | TCATAAAATTGATAAGGAGAATCC | Amplification of the URA3 gene plus 1Kb homologies |
| URA3KI1000_R | GTTATCAGATATTATCAGGTGG | Amplification of the URA3 gene plus 1Kb homologies |
| URAKOver_F | CTGCCAAGCTATTTAATATCA | Confirmation of URA3 gene KO |
| URAKOKIver_R | AAGTAACAGTATTTTACGGGG | Confirmation of URA3 gene KO and KI |
| URAKIver_F | AAAAGGATTAAAGATGCTAAGAG | Confirmation of URA3 gene KI |
| LEU_F | GATCGTTTTGTTAGGTGCTGTGGG | Hybridization of a dsOligo contaning a sgRNA sequence targeting LEU2 |
| LEU_R | AAACCCCACAGCACCTAACAAAAC | Hybridization of a dsOligo contaning a sgRNA sequence targeting LEU2 |
| XR_F | GTTTCGACGGATTCTAGAACTAGTGGATCCATG CCTTCTATTAAGTTG | Amplification of Sc. stipitis xylose reductase (xyl1) |
| XR_R | TCGAATTCCTGCAGCCCGGGGGATCCTTAGACG AAGATAGGAATC | Amplification of Sc. stipitis xylose reductase (xyl1) |
| P425_F | GCTGAAATATACGGGTTCCC | Amplification of the XR expressing cassette in p425 |
| P425_R | CTGGTAGGTTAGATCCCAGG | Amplification of the XR expressing cassette in p425 |
| 29_F | GATCCTTACATGTTTGGCACGCAG | Hybridization of a dsOligo contaning a sgRNA sequence for pGS.29 |
| 29_R | AAACCTGCGTGCCAAACATGTAAG | Hybridization of a dsOligo contaning a sgRNA sequence for pGS.29 |
| 30_F | GATCCTTACATGTTTGGCACGTAG | Hybridization of a dsOligo contaning a sgRNA sequence for pGS.29 |
| 30_R | AAACCTACGTGCCAAACATGTAAG | Hybridization of a dsOligo contaning a sgRNA sequence for pGS.30 |
